# High-resolution protein fragment interactions using AVA-Seq on a human reference set

**DOI:** 10.1101/2021.07.28.454266

**Authors:** Stephanie Schaefer-Ramadan, Jovana Aleksic, Nayra M. Al-Thani, Yasmin A. Mohamoud, David E. Hill, Joel A. Malek

## Abstract

Protein-protein interactions (PPIs) are important in understanding numerous aspects of protein function. Here, the recently developed all-vs-all sequencing (AVA-Seq) approach to determine protein-protein interactions was tested on a gold-standard human protein interaction set (hsPRS-v2). Initially, these data were interpreted strictly from a binary PPI perspective to compare AVA-Seq to other binary PPI methods tested on the same hsPRS-v2. AVA-Seq recovered 20 of 47 (43%) binary PPIs from this reference set comparing favorably with other methods. The same experimental data allowed for the determination of >500 known and novel PPIs including interactions between wildtype fragments of tumor protein p53 and minichromosomal maintenance complex proteins 2, and 5 (MCM2 and MCM5) that could be of interest in human disease. Additional results gave a better understanding of why interactions might be missed using AVA-Seq and aide future PPI experimental design for maximum recovery of information.

## Introduction

Understanding protein-protein interactions (PPIs) by uncovering interacting regions and/or active sites has been important for the advancement of many biological fields. Knowing a protein’s partners allows for innovation of drug discovery by being able to interrogate active sites and interaction interfaces for clinically relevant inhibitors. When researchers can connect or extend protein interaction networks new information can be utilized to predict function of unknown genes.

The yeast two-hybrid (Y2H) method revolutionized how interacting partners could be determined^1^, opening the way for systematic, proteome scale binary interaction mapping for human and model organisms ^2–8^. Many advancements in binary interaction mapping since then have added to the conversation with no single method being superior to another - meaning no one method can determine all or most PPIs without systematic bias. A recent manuscript^9^ illustrates the complexities of the PPI process by utilizing a human positive reference set (hsPRS-v2) which contained 60 human interacting protein pairs. At best, the use of one method in isolation could determine 33% of the hsPRS-v2 and using 10 versions of 4 assays could recover 63% of interactions^9^. Importantly, Choi and colleagues confirm the significance that assay configuration and orientation has on interaction screening. Meaning fusions to different individual DNA binding (DB) domains or different transcriptional activation domains (AD) to reconstitute transcription factor activity can have non-overlapping results or unforeseen bias making the screening area at least 2-fold higher to ensure proper coverage of the interaction space. To achieve maximal detection of binary interactions, multiple methods that incorporate different assay configurations and fusion partner orientations will need to be employed to gain significant coverage of the interactome in question.

The all-vs-all sequencing (AVA-Seq) method is based on a bacterial two-hybrid system and with the innovation of a plasmid which allows convergent fusion proteins was recently developed^10^. This system allows the use of next-generation sequencing (NGS) to determine protein interactions on a large scale as well as providing a level of domain-domain interaction information due to the testing of multiple overlapping protein fragments present in the library. While results from an initial test on 6 human proteins were encouraging, it is important to put the system in the context of other methods by using a gold-standard set of interactions. To that end, the AVA-Seq system was applied on a subset of the hsPRS-v2 proteins, a gold-standard set of PPIs. The hsPRS-v2 was used in this study as a validation tool to test the sensitivity of the AVA-Seq method. Current methods in the field are limited by the ability to scale, high cost, and laborious colony selection. AVA-Seq is a novel way to screen PPI in a fast, cost-effective way and was designed to screen fragmented proteins against itself (or an alternative library) incorporating high sensitivity and multiple orientations simultaneously.

## Results

### Overview of AVA-Seq Method

The hsPRS-v2^9^ was utilized to compare the ability of AVA-Seq to recover binary interactions with other methods, with the exception that a library of protein fragments for each hsPRS-v2 protein was tested as opposed to simply full-length proteins. Fig. 1 illustrates the method approach utilized for this study. First, two separate pools of positive reference set (PRS) proteins were made from the hsPRS-v2 library (Supplemental Table 1). ‘PRS Batch 1’ contained 39 PRS proteins and 9 random reference set (RRS) proteins believed to not interact while ‘PRS Batch 2’ contained 41 PRS proteins and 9 RRS proteins. Both batches were prepared as separate experiments (meaning no cross interactions between batches would be detected) but processed in parallel. Briefly and as described in Andrews et al.^10^ the specific proteins for each batch (Supplementary Table 1), including selected RRS proteins, were pooled separately, sheared, size selected and ligated into pBORF-AD and pBORF-DBD. After selection for open reading frames (ORFs), fragments were amplified, ‘stitched’ together using overlap extension PCR and ligated into pAVA for screening. Screening consisted of triplicate samples grown under varying selective media (0 mM, 2 mM and 5 mM 3-AT). Then the surviving plasmids were sequenced using NGS to detect differential growth among the various conditions. Two separate transformation and screening events were conducted for each PRS Batch (i.e., Batch 1A and 1B and should be considered biological replicates as the plasmids came from the same DNA pool but were transformed separately). Data analysis for PRS Batch 1 and 2 were performed identically but separately since the protein pools are unique (Fig. 1B). For each batch, the expected binary interactions for hsPRS-v2 were determined (Supplemental Table 2) and a cumulative table of all-vs-all interactions (PRS Batch 1 and 2 combined) were populated (Fig. 1B, Supplemental Table 3).

**Fig. 1:**
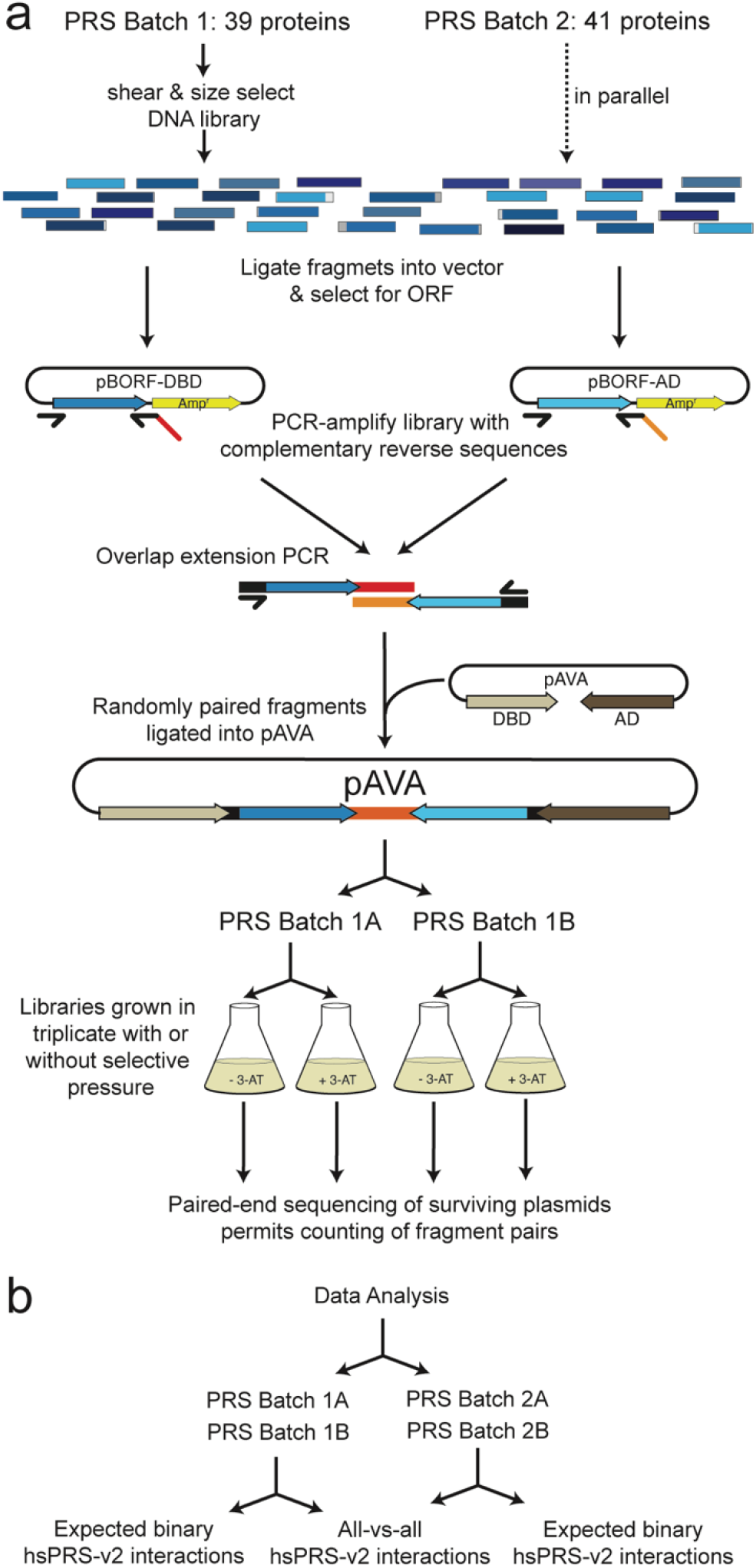
Method schematic. **a**) PRS batch 1 (39 proteins) and batch 2 (41 proteins) were treated as separate experiments and processed in parallel (Supplemental Table 1). First, the proteins were pooled, sheared, size selected and ligated into pBORF-AD and pBORF-DBD. After selection for the open reading frame (ORF), fragments were amplified, ‘stitched’ together using overlap extension PCR and ligated into pAVA for screening. For each PRS batch two separate screenings (A and B) were conducted, and the data generated were pooled during analysis. **b**) Data analysis for Batch 1 and 2 were performed identically but separately since the protein pools are unique. For each batch, the expected binary interactions were determined (Table 1 and Supplemental Table 2) and a cumulative table of all-vs-all interactions (Batch 1 and 2) were populated (Supplemental Table 3). Batch 1 and 2 included an additional 9 RRS proteins for control.

An important aspect of AVA-Seq, especially when the interaction pools are relatively small, is the open reading frame (ORF) filtering. This step is significant because as the protein pools being tested become larger the screening area also increases by a factor of 36 (6 by 6 possible reading frames) making the likelihood of both fragments being in frame 1 to be 2.7% without ORF filtering. With this study, nearly 80% of the fragments associated with DBD and AD have been enriched for frame 1 (data not shown). After “stitching” the DBD and AD fragments together, 64% of convergently fused fragments generated were in frame 1. ORF filtering readily allowed greater than 3-fold coverage of the interaction space in a short amount of time without exhausting resources. One benefit of using fragments over full-length proteins is in the context of an auto-activating protein. Meaning, not all fragments from a protein might auto-activate the system by interacting with RNAP (activation domain; AD) or lambda cI (DNA binding domain; DBD). Therefore, only the in-frame fragments that interact with multiple out-of-frame fragments needs to be removed as these are suspected to be possible examples of auto-activation^11,12^. Here, 13 fragments that were fused to RNAP and interacted with more than 3 out of frame fragments fused to lambda cI were removed. These are suspected to actually interact with lambda cI and auto-activate. Similarly, 21 fragments were removed that auto-activate by interaction with RNAP. These analyses were only conducted on 2 mM conditions as the 5 mM conditions did not show signs of significant numbers of auto-activators.

### Analysis of sequence data

While the concept of deep sequencing to identify a difference in counts of fragments in libraries tested in various conditions has been used in multiple ways, some adjustments were required from the standard analyses designed for techniques such as RNA-seq. Detailed investigation of the data set consistently showed a decrease of counts values of many fragment pairs from 0 mM to 2 mM 3-AT, and smaller decrease from 0 mM to 5 mM 3-AT. Potential interaction of thousands of fragment pairs in 2 mM 3-AT, the less selective condition, takes a larger portion of the read counts, causing the non-interacting fragment pairs to decrease in overall proportion. Under standard assumptions this would result in a “negative” interaction being observed, i.e. a decrease in sequences from a protein pair under selective conditions. To address this, the data were scaled appropriately based on multiple factors as discussed (see Methods).

For each protein pair tested, the percentage of the total possible test space covered by at least one fragment from each protein was documented and plotted in Fig. 2. As explained in the method overview section above, the protein interactions were split into two separate batches. Orientation of the fragment pairings with respect to the activation domain (AD) or the DNA binding domain (DBD) are illustrated in Fig. 2. 73.2% and 69.7% of the total possible search space were covered by at least one in-frame fragment for both Batch 1 and 2, respectively. While the total percent of the possible search space covered by at least one in-frame fragment is high, it is apparent there are proteins which have coverage in one orientation but not the other (i.e., IFG2 and MAFG in PRS Batch 1; Fig. 2). Moreover, it was observed that ORF filtering yielded poor coverage or complete absence of proteins which are < ~300 amino acids in length. Indeed, only 25% of interactions involving one protein of 300 amino acids or less were recovered. In this study, the average full-length of interacting proteins were 534 amino acids. The average full-length protein for interactions that were not detected in this study but had at least one protein fragment pair with the expected interacting partner were 339 amino acids. This is likely a result of the fact that fragments were size selected for approximately 450 base pairs (150 amino acids) and that may reduce the chances of capturing multiple fragments without stop codons when ORF filtering is applied. Yet, for proteins with a length greater than 300 amino acids 75% of interactions were captured. This argues for the sensitivity of the AVA-Seq system when the conditions are right. Specifically, coverage and depth of coverage of a protein by tested fragments are important and likely affected by length when ORF filtering is used. Protein pairs with at least one short protein were less likely to have a known interaction detected in the system (Fig. 3a). That is, when the expected interacting proteins have full-lengths > 434.7 amino acids, on average, there is a significantly higher chance of detecting the interaction when compared to expected interacting pairs which did not show an interaction (mean full-length of 245.7 amino acids) (p-value=4.40411×10^−05^). Likewise, as the number of unique fragments which represent the expected interaction increase relative to minimum protein length (depth of coverage) there is a statistically significant increase to detect the interaction (p-value = 9.0144×10^−07^) (Fig. 3b). The idea of improvement in detecting an interaction by deeper coverage is not due to simply increased chances of detecting a random interaction is discussed below. To investigate whether the bias against detecting interactions from pairs with at least one shorter full-length protein was not due to an inherent bias of the system against shorter proteins the full-length of the protein was plotted vs the depth of coverage by fragments (Fig. 3c). As Figs. 3a and b suggest, it is likely that interactions were not detected simply because shorter proteins were less likely to have sufficient depth of coverage. That is, they might have a single fragment covering the majority of the length of the protein (Fig. 2a and b) but multiple fragments appear to be beneficial in detecting an interaction and these only increased with the length of the protein.

**Fig. 2:**
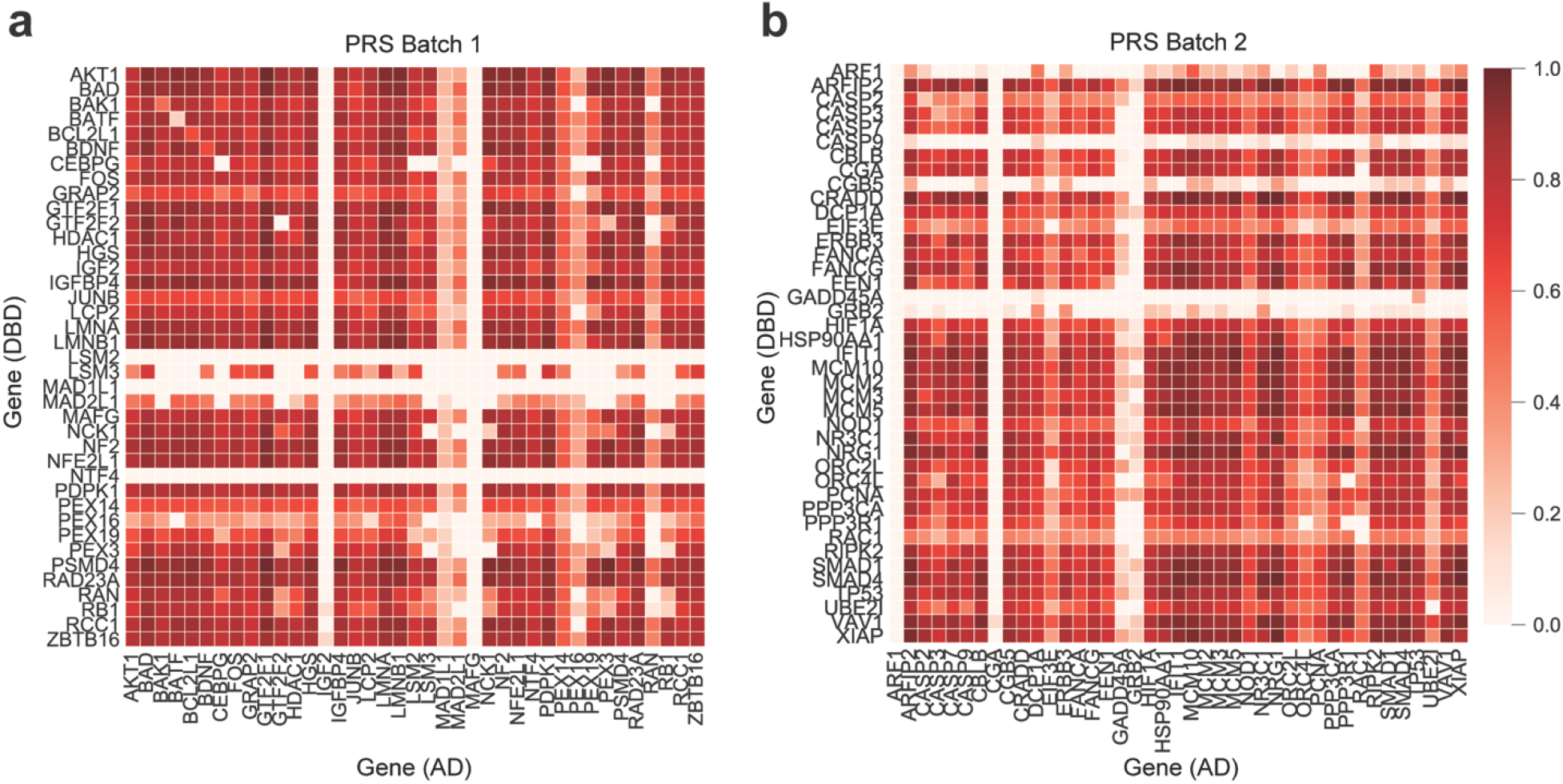
Heat maps of gene coverage. **a**) positive reference set (PRS) Batch 1 (39×39 proteins) and **b**) PRS Batch 2 (41×41proteins). Color scale indicates percent gene coverage in a specific orientation (AD or DBD associated) with 1 being 100% coverage of the protein and 0 representing 0% coverage. RRS proteins are not included.

**Fig. 3:**
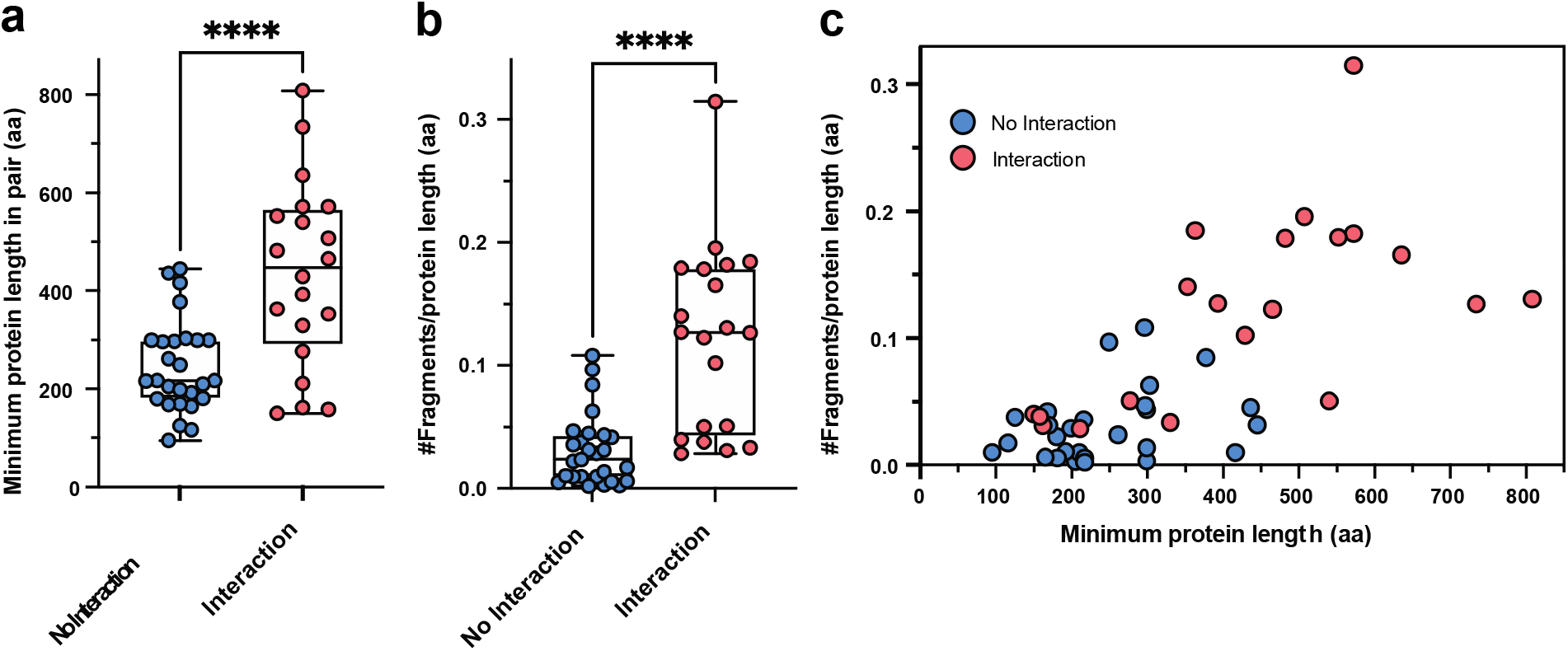
Protein length vs interaction. **a**. individual protein length in amino acids of proteins used in this study categorized into no interaction (blue; mean 245.7; n = 27) or interaction observed (red; mean 434.7; n = 20; t=4.524, df=45). P value <0.0001 indicated. **b**. the minimum number of relative fragment starting points divided by protein length in amino acids vs. no interaction (blue; mean 0.03088; n = 27) or interaction observed (red; mean 0.1211; n = 20; t=5.689; df=45). P value <0.0001 indicated. **c**. The number of protein fragments per protein length (in amino acids) plotted against the minimum protein length in the expected interacting pair. Blue dots represent no interaction and Red dots represent interaction observed.

### Binary Interactions

AVA-Seq recovered 20 of the 47 (42.55%) PPIs tested from the hsPRS-v2 (Table 1). Of the 20 binary interactions detected using AVA-Seq a few hundred fragments passed filtering whereas a few thousand did not showing the selectivity of the system. Of the 20 hPRS-v2 pairs that AVA-Seq detected as domain-domain interactions, five were not captured by other assays^9^ (Table 1). Additionally, at least two phosphorylation dependent protein interactions (TP53:UBE2I and SMAD1:SMAD4) were recovered highlighting the ability to identify potentially novel interaction regions between proteins which typically require a post-translational modification and are not feasible to detect with a bacteria system. Why this is possible is not yet clear but will be of interest in future investigations.

**Table 1.** Detected binary PPI recovered from hsPRS-v2. The protein interaction pair numbering in the left most column as well as protein naming is according to Choi, et al.^9^. The asterisk (*) denotes the PPI was recovered using a method published in Choi, et al. The last five without an asterisk are interactions which were recovered uniquely by AVA-Seq.

|  | Protein1 | Protein2 | Recovered by other methods<sup>9</sup> |
| --- | --- | --- | --- |
| 2 | LMNA | LMNB1 | * |
| 5 | LCP2 | GRAP2 | * |
| 6 | BAK1 | BCL2L1 | * |
| 11 | PSMD4 | RAD23A | * |
| 15 | MAFG | NFE2L1 | * |
| 27 | MCM2 | MCM3 | * |
| 28 | AKT1 | PDPK1 | * |
| 30 | NF2 | HGS | * |
| 31 | TP53 | UBE2I | * |
| 32 | HIF1A | TP53 | * |
| 34 | SMAD1 | SMAD4 | * |
| 35 | CEBPG | FOS | * |
| 37 | SMAD4 | DCP1A | * |
| 40 | NR3C1 | HSP90AA1 | * |
| 46 | LMNA | RB1 | * |
| 49 | ORC2L | MCM10 |  |
| 51 | HDAC1 | ZBTB16 |  |
| 52 | XIAP | CASP3 |  |
| 55 | RIPK2 | NOD1 |  |
| 59 | MCM2 | MCM5 |  |

As mentioned above, there was a clear trend for detecting interactions where one of the interacting partners was >350 amino acids. While the average full-length protein in the study was 534 amino acids, in the case of the 27 binary interactions that were not detected, 23 (85%) contained one partner with a full-length less than 350 amino acids.

### Sensitivity and Selectivity of AVA-Seq System

The sensitivity of AVA-Seq was controlled on a basic level by the addition of a known a protein interacting pair in the pAVA vector (LGF2-Gal11p). This control was added to each library at the screening stage (see Methods) and showed consistently strong results in both 2 mM and 5 mM conditions with and average logFC and average FDR of 7.09 and 1.76×10^−13^, respectively.

The selectivity of a particular system is put to the test when all permutations are considered such as in an all-vs-all screen. Under these conditions, potentially millions of pair-wise interactions are tested and the chance for significant numbers of random interactions increases unless the correct selection criteria are applied. In some cases, single proteins were represented by hundreds of fragments that were screened against thousands of fragments from all other proteins. The basic statistical cutoffs using a log2 fold change (LogFC) of 1 and false discovery rate (FDR) of 0.1 are selective and resulted in 2,606 unique fragment pairs called as interacting from a total of 283,676 non-interacting fragments. That is, 0.91% of the total fragments that were considered for statistical testing were involved in a possible interaction.

Furthermore, the data were searched for evidence that fragments called as interacting were not simply random representations of all screened fragments. Multiple approaches were employed to find evidence of interacting fragments which overlapped known interaction domains, interacting fragments which lay outside of non-interacting fragment peaks, and interacting fragments which were more localized and less disperse than random fragments as would be expected if the interacting fragments were covering a true interacting domain. First, evidence for fragments identified as interacting in the system and overlapped with previously identified interacting regions were identified. As an example, fragments called as significant in the system for the HGS:NF2 interaction were plotted (Fig. 4). In both the HGS and NF2 examples the fragments that were enriched under selective pressure indicate an interaction (red trace) and align well to the interacting regions from the literature^13–15^ (grey shaded box(es) in Fig. 4a and 4b). This is remarkable especially given the thousands of protein fragment pairs which did not pass filtering criteria as an interaction, indicating a highly selective screening method. Indeed, there were regions of proteins with very high counts of fragments paired with other proteins, but which did not yield any called as interacting confirming that it is not simply random fragment pairs that pass the filtering criteria. As a follow up and to demonstrate that interacting protein fragments are not simply randomly drawn from non-interacting fragments the average gap between fragment start points for non-interacting fragments were compared to those of interacting fragments. The goal was to show the localization of interacting fragments is not random across the protein but more likely to be localized assuming there is one interacting domain. Fig. 4e plots full-length protein >450 amino acids (~3x average fragment size) vs average distance between fragments in amino acids. Unique interacting fragment pairs (where at least one of the fragments start points were different) were extracted from the all-vs-all data and distance between their start points averaged. The data for non-interacting fragments were generated using random starting point picking, 1,000 times. The R^2^ values for the random and interacting starting points were 0.9917 and 0.4961, respectively. The poor linear fit of the interacting fragment start points is additional evidence these not random screening events. Fig. 4f plots the paired t-test indicating a statistically significant p-value <0.0001.

**Fig. 4:**
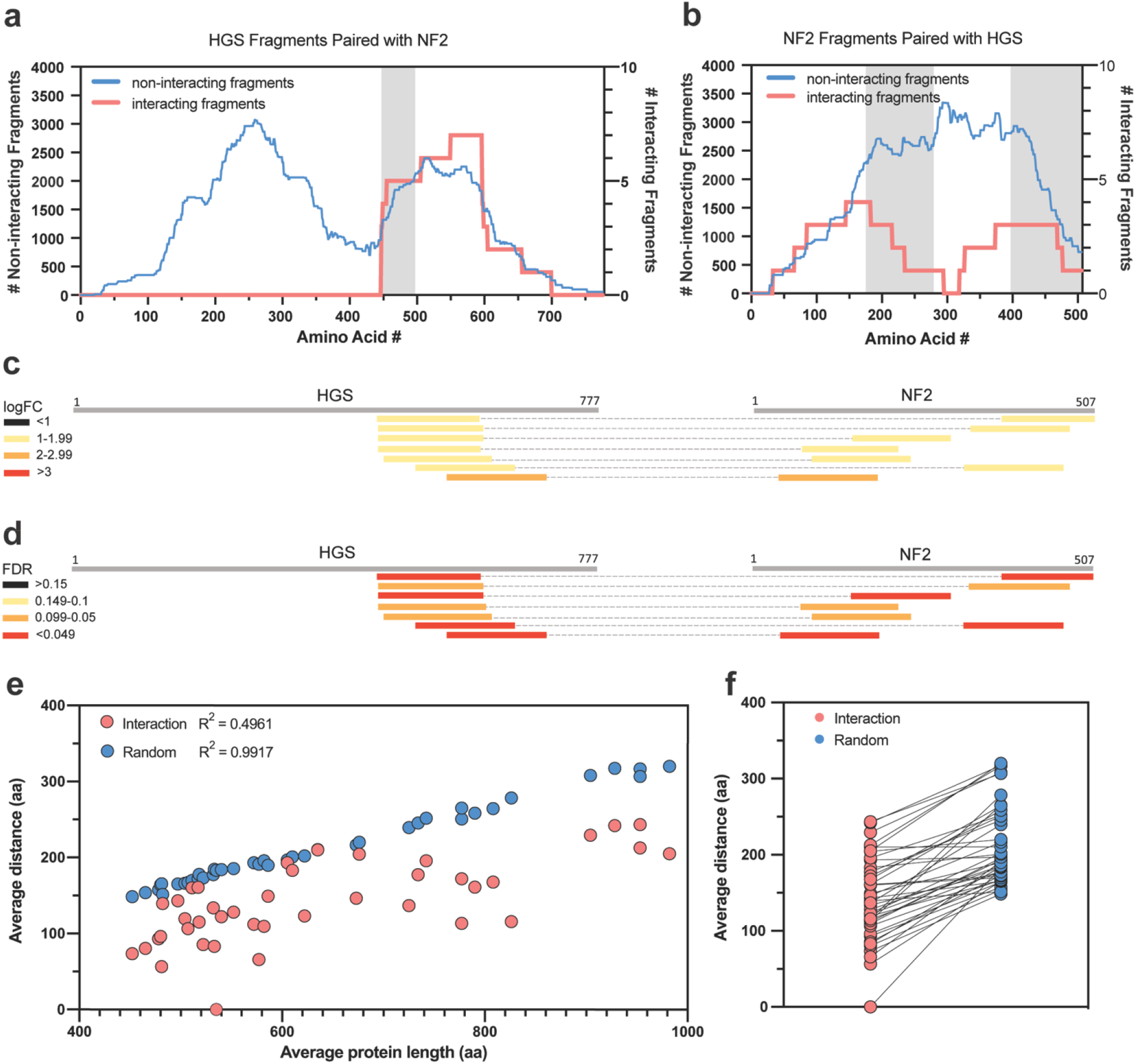
Selectivity of fragment interaction. Panels **a** and **b** illustrate the selectivity of the interacting fragments between HGS and NF2 genes. The blue traces (**a** and **b**) represent non-interacting fragments (left y axis) vs fragment start point while the red traces (**a** and **b**) represent interacting fragments (right y axis) vs fragment start point. The grey shaded regions in **a** and **b** highlight the expected interaction region of HGS with NF2 from the literature^14^. Panels **c** and **d** illustrate the fragment pairings between HGS and NF2 along with logFC and FDR, respectively. **e**. the average fragment distance in amino acids (aa) plotted against the average protein length. Protein fragments utilized in this plot were associated with proteins that had at least two interacting start points fragments with at least one other interacting partner. The average distance of interacting starting points was then computed. **f**. paired t-test for data in panel. (t=10.84; df=40).

The tested set included randomly selected protein pairs (RRS) for which no evidence of an interaction has been reported^9^ and the threshold for determining the percentage PRS detection is set at a zero RRS detection of pairs of full-length proteins^9^. Six of 12 RRS protein pairs did not interact in our system and the remaining 6 showed statistical significance using the same criteria as the binary interactions (logFC >1, FDR <0.1) (Supplementary Figs. 1–2). Five of the 6 RRS interacting pairs had more than one fragment pair and one pair survived the stringent 5 mM 3-AT growth conditions which would indicate a strong interaction in our system. These interactions need to be investigated further to understand if they are indeed biologically relevant or if domains interacted due to being surface exposed as a result of the fragment approach used here.

### Considering the RRS interactions in the context of AVA-Seq

Six pairs of RRS (random reference set) protein interactions were included in each PRS batch (Supplementary Table 1). Normally, the RRS interactions are used to calibrate the assay allowing researchers to directly compare protein-protein interaction methods^9^. The RRS results are used as a threshold, meaning that any interaction detected in the PRS be stronger (above the threshold) when compared to the strongest RRS interaction. More stringent criteria were applied to these data with the intention to reduce the RRS interactions significantly while still maintaining a reasonable number of PRS interactions. Keeping in mind the reduction of RRS interactions also means more real interactions are being lost.

The table of 584 all-vs-all interactions has 5 RRS interactions and 16 PRS interactions which were recovered (Supplemental Table 2; Supplemental Fig. 4). Upon applying even more stringent criteria, 584 interactions reduce to 431 and subsequently the PRS interactions reduce from 16 to 12 (out of 47) and the RRS interactions reduce from 5 to 2 (out of 12) (Supplemental Fig. 4). The more stringent filters are effectively requiring “stronger” interactions. That is, they either passed in the 5 mM 3-AT conditions or had even higher fold change in the 2 mM conditions. The fact that 2 of the RRS pass this means to us that, while possibly not biologically relevant, they are reproducibly interacting in the system. If the number of RRS is reduced to 0 simply on the criteria of “stronger interaction”, the resulting data might not always be a proxy for a biologically relevant interaction. With the knowledge that interaction strength/reproducibility in an *in-vitro* system may not automatically equate to biological relevance we recommend that future studies provide tables of reasonable cutoffs but also all data so each investigator can make decisions on thresholds appropriately. That’s a strength of the various levels of interaction quality information obtained from AVA-Seq.

We realize our method is a nontraditional use of the PRS and RRS and this makes it difficult to compare directly to other studies of the same gold standard interaction set. Specifically, our use of fragments over full-length proteins may cause interactions (both biologically relevant and irrelevant) to be detected. Therefore the ‘answer’ for this manuscript would be different than other binary applications. Especially given we are not able to calibrate our data in the same manner as demonstrated previously^9^ due to the use of fragments as opposed to strictly full-length proteins. However, we are confident that our system is detecting interactions reproducibly across a range of strengths and believe the user can filter the data based on what their needs may be. Additional measures of stringency could also be applied such as increasing 3-AT concentrations are using multiple reporters to generate an interaction score or confidence score^16^.

### All-vs-All Analysis on data set meant for Binary tests

The original intent of this study was not to uncover novel interactions among all hsPRS-v2 proteins, however, due to the inherent design of AVA-Seq these data were automatically populated. In the context of an all-vs-all analysis of the data the search space of interactions is dramatically increased, and therefore more stringent criteria needed to be applied. A benefit of the AVA-Seq system includes fragments being used in multiple fusion orientations and inhibitor concentrations allowing for multiple independent tests of the same interaction. An inexhaustive list of known PPIs were identified using this approach (Table 2) as well as many novel interactions (selected interactions listed in Table 3). These interactions are supported by multiple criterion which gives great confidence to the robustness of the interaction data, at least in the context of this *in vitro* screen. These criteria include fragments with similar start positions, fragments being both AD and DBD associated, interactions detected under both 2 mM and 5 mM 3-AT selective pressure and interactions being detected in multiple unique transformation events. A total of 901 PPIs were detected with any form of evidence, however applying simple criteria such as requiring multiple fragments or fragments in both orientations (see Methods) reduced this to 584 interactions among the PRS and RRS proteins (Supplemental Fig. 4). The same criteria applied to the binary interactions reduced those from 20 detected interactions to 16 (a 20% loss of known interactions). The 901 PPIs detected in the all-vs-all data is 37%, while the 584 PPIs are 24% of 2451 possible interactions given the batch sizes of 48 and 50 proteins (including RRS proteins; total possible interactions calculated using n*(n+1)/2). Other studies conducted with AVA-Seq on randomly selected proteins have shown the expected scale-free nature of the proteins with most proteins having few interacting partners (unpublished data). However, in this study, we did not observe a scale-free network and observed steady decrease in interacting partners (Supplementary Figure 3) indicating this may be due to the non-random selection of the PRS proteins.

**Table 2.** Known interactions detected using all-vs-all. Selected known interactions from the combined all-vs-all data from Batch 1 and Batch 2 including RRS proteins. This list is a subset of Supplemental Table 3.

| Previously known Protein-Protein Interactions confirmed in this study |  |  |  |  |  |  |  |  |  |  |
| --- | --- | --- | --- | --- | --- | --- | --- | --- | --- | --- |
| Protein1 | Protein2 | Orient1 | Orient2 | 2 mM | 5 mM | logFCmax | FDRmin | #Libs | #uniqFragPairs | Reference |
| MCM5 | MCM3 | 31 | 5 | 19 | 17 | 6.47421451 | 3.12E-09 | 2 | 29 | 22,23 |
| MCM2 | MCM10 | 5 | 7 | 10 | 2 | 3.62060897 | 7.00E-05 | 2 | 11 | 24 |
| MCM2 | MCM3 | 56 | 5 | 38 | 23 | 3.0538095 | 1.64E-13 | 2 | 46 | 22,24 |
| ORC2L | MCM3 | 2 | 5 | 5 | 2 | 3.69014656 | 0.00897978 | 1 | 7 | 25,26 |
| FOS | LMNA | 14 | 7 | 19 | 2 | 6.34040694 | 2.21E-235 | 1 | 15 | 27 |
| HSP90AA1 | HSP90AA1 | 4 | 4 | 6 | 2 | 9.03727216 | 1.25E-12 | 1 | 6 | 28 |
| HSP90AA1 | TP53 | 2 | 2 | 4 | 0 | 3.54728603 | 2.84E-29 | 2 | 4 | 28 |
| CASP3 | XIAP | 2 | 0 | 1 | 1 | 4.71341933 | 6.82E-21 | 1 | 1 | 29-31 |
| XIAP | RIPK2 | 1 | 0 | 1 | 0 | 2.62693776 | 0.07813117 | 1 | 1 | 3,32 |
| RIPK2 | Nod1 | 1 | 0 | 1 | 0 | 2.02393071 | 0.06723463 | 1 | 1 | 33 |
| NOD1 | HSP90AA1 | 5 | 0 | 4 | 1 | 5.39888581 | 0.01624673 | 1 | 4 | 34 |
| NOD1 | XIAP | 3 | 0 | 3 | 0 | 3.15614296 | 0.01775524 | 1 | 3 | 32 |

**Table 3.**
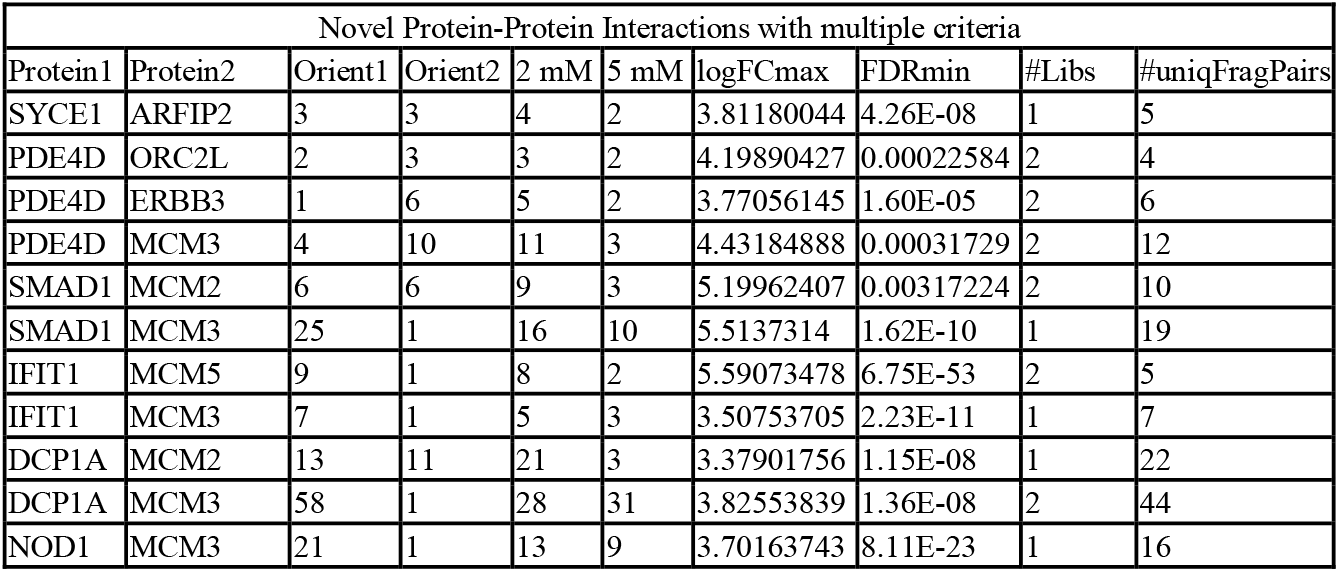
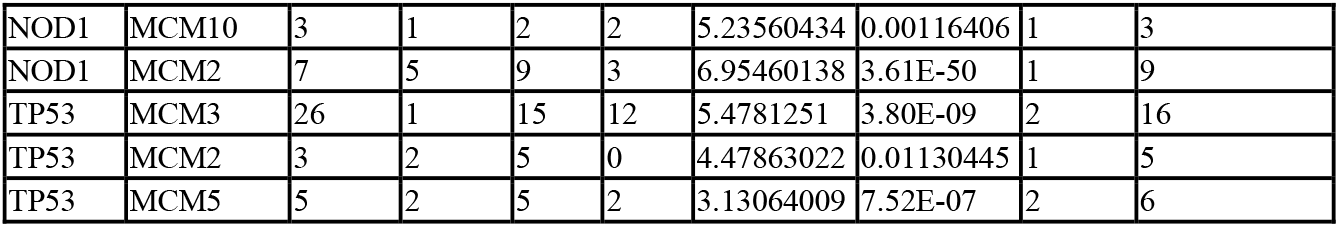
Novel interactions detected using all-vs-all. Selected novel interactions from the combined all-vs-all data from Batch 1 and Batch 2 including RRS proteins. This list is a subset of Supplemental Table 3.

| Novel Protein-Protein Interactions with multiple criteria |  |  |  |  |  |  |  |  |  |
| --- | --- | --- | --- | --- | --- | --- | --- | --- | --- |
| Protein1 | Protein2 | Orient1 | Orient2 | 2 mM | 5 mM | logFCmax | FDRmin | #Libs | #uniqFragPairs |
| SYCE1 | ARFIP2 | 3 | 3 | 4 | 2 | 3.81180044 | 4.26E-08 | 1 | 5 |
| PDE4D | ORC2L | 2 | 3 | 3 | 2 | 4.19890427 | 0.00022584 | 2 | 4 |
| PDE4D | ERBB3 | 1 | 6 | 5 | 2 | 3.77056145 | 1.60E-05 | 2 | 6 |
| PDE4D | MCM3 | 4 | 10 | 11 | 3 | 4.43184888 | 0.00031729 | 2 | 12 |
| SMAD1 | MCM2 | 6 | 6 | 9 | 3 | 5.19962407 | 0.00317224 | 2 | 10 |
| SMAD1 | MCM3 | 25 | 1 | 16 | 10 | 5.5137314 | 1.62E-10 | 1 | 19 |
| IFIT1 | MCM5 | 9 | 1 | 8 | 2 | 5.59073478 | 6.75E-53 | 2 | 5 |
| IFIT1 | MCM3 | 7 | 1 | 5 | 3 | 3.50753705 | 2.23E-11 | 1 | 7 |
| DCP1A | MCM2 | 13 | 11 | 21 | 3 | 3.37901756 | 1.15E-08 | 1 | 22 |
| DCP1A | MCM3 | 58 | 1 | 28 | 31 | 3.82553839 | 1.36E-08 | 2 | 44 |
| NOD1 | MCM3 | 21 | 1 | 13 | 9 | 3.70163743 | 8.11E-23 | 1 | 16 |
| NOD1 | MCM10 | 3 | 1 | 2 | 2 | 5.23560434 | 0.00116406 | 1 | 3 |
| NOD1 | MCM2 | 7 | 5 | 9 | 3 | 6.95460138 | 3.61E-50 | 1 | 9 |
| TP53 | MCM3 | 26 | 1 | 15 | 12 | 5.4781251 | 3.80E-09 | 2 | 16 |
| TP53 | MCM2 | 3 | 2 | 5 | 0 | 4.47863022 | 0.01130445 | 1 | 5 |
| TP53 | MCM5 | 5 | 2 | 5 | 2 | 3.13064009 | 7.52E-07 | 2 | 6 |

## Discussion

The AVA-Seq system takes advantage of NGS to significantly increase either the breadth or resolution of a PPI screen providing evidence for domain-domain level interaction information. Using a gold standard protein reference set, AVA-Seq recovered 20 of 47 PPIs with 5 (25%) of these binary interactions being unique to the AVA-Seq method^9^. It is very likely the assay properties of AVA-Seq enrich for PPIs which would not normally be ‘detectable’ using existing two-hybrid assays, particularly those relying on expression of full-length proteins^17^. This is likely due to the fusion of smaller protein fragments which are either easier to express or are more exposed relative to a full-length protein. A small percentage of the human interactome is comprised of very stable and functionally conserved interactions^4^. Because AVA-Seq was able to recover unique interactions, it is possible this method has an advantage for screening intrinsically disordered proteins. Even though the intention of this study was to compare the ability of AVA-Seq to determine known human PPIs with other binary methods, AVA-Seq has additional advantages such as populating protein fragments in an all-vs-all fashion. AVA-Seq has dual orientation fusions built into the design. This aspect alone should increase detection sensitivity by at least 1.3-fold within a single assay^9^. Additionally, Choi and colleagues expanded on the idea that permuting the experimental conditions has added benefit. Using 10 versions of 4 assays Choi et al. demonstrated 63% recovery of PPIs using hsPRS-v2 as a standard^9^. Since AVA-Seq uses fragmented proteins rather than full-length proteins, having a PPI requirement to have more than one fragment start point and appear in both orientations has significant added value when determining novel interactions or increasing the resolution of a protein interaction site.

For the all-vs-all data, a large fraction of all possible combinations, 37% with minimal filtering and 24% with expanded filtering, were recovered as having some evidence for interacting. While this fraction is high with respect to other studies, it is important to note that the proteins used here are not randomly selected but may be more biased towards proteins that interact with many partners. Typically, proteins with a connectivity above a certain threshold are removed but that was not possible here given the steady trend of decrease in connectivity. Rather than interpreting this as a lack of specificity we consider that the non-random selection of the PRS proteins may contribute but further investigation will be necessary.

In the novel set of interactions, several were of interest to human disease. Specifically, the TP53:MCM5^18^ and TP53:MCM2^19^ proteins have been associated previously. Interestingly, both were associated with TP53 gain-of-function mutations and, at least in the case of the MCM2 interaction, wild-type interactions were not consistently detected. It is possible that the gain-of-function mutations increases the strength of the interaction to a level that *in vitro* systems could detect even though the interaction would not be observed under wild-type conditions; however, the AVA-Seq method, utilizing small protein fragments with multiple start and stop fusions, was able to detect the interactions. While the TP53 used in this study contains P72R and P278A point mutations not all fragments necessarily contain these mutations. While a few of the fragment pairs between TP53:MCM5 and TP53:MCM2 did include a P278A mutation which is part of the hsPRS-v2 template for TP53, significant interactions were also detected with TP53 fragment which did not include this portion of the protein sequence. There were no noticeable differences between fragment pairs containing or lacking the P278A mutation in terms of strength of the interaction. While interactions were detected between fragments of wild-type TP53:MCM5 even in the more stringent 5 mM 3-AT conditions and TP53:MCM2 had interactions only in the 2 mM condition. These interacting pairs hint that interactions between wildtype TP53 and MCM proteins are likely. A future study could utilize the AVA-Seq system to look at gain-of-function mutations versus wild-type to see if the mutation(s) does indeed simply increase the strength of the interaction rather than create it *de-novo*.

As with any method there are limitations which exist, and they become clearer as different data sets are applied and different scientific questions are being asked. There are instances where there is an indicated interaction in 5 mM but not in 2 mM selection media despite 2 mM being the less selective condition. It is possible that deeper sequencing of 2 mM replicates when compared to 5 mM replicates may be necessary as there are significantly more interactions which occur under the less stringent 2 mM conditions. Because of this, the question remains how ‘deep’ of sequencing is needed for the 2 mM replicates. It is clear from these data that the more unique fragments a protein has increases the chance of detecting an interaction. This notion helps reiterate that more fragments overlapping a given area allows to not only increase chances of detecting an interaction but increase resolution of the protein interaction region with a given protein or set of proteins. Notably, these are not just a function of increased random fragments being detected as interactions and indeed remains selective is discussed below. Another interesting question uncovered was regarding the feasibility of ORF filtering with short proteins. As indicated in Fig. 3, there is a significantly higher chance to detect a protein interaction if both proteins are longer because of increased probability that ORF selection produces more overlapping fragments for those proteins. There are several potential ways to mitigate these affects in future studies. First, for more focused protein network studies, such as this, smaller shearing (i.e., 250-300 base pair instead of 450 base pair) with no ORF filtering would allow for smaller proteins to make it into the final fragment pool and eliminate one source of bias. The benefit of this system is tested fragments are C-terminal to the fusion proteins allowing the testing of fragments that include stop codons. Another option would be to synthesize gene fragments of the proteins eliminating the need for ORF filtering. Although the ORF filtering is essential to reduce the screening area when screening large protein pools (Schaefer-Ramadan, unpublished), there may be significant value in terms of interaction resolution in generating protein fragment libraries which have not been subjected to ORF filtering in addition to offering a higher depth of fragment coverage. It is worth noting that previous work identified different populations of interacting fragment start points when comparing ORF filtered fragments to those which were not^10^. Limitations exist regarding the fragment length amendable to NGS technology. 850-900 base pair libraries can consistently be paired-end sequenced using Illumina technology limiting individual fragments to approximately 450 base pair. As with any bacterial system used to express human proteins, interactions requiring one or more post-translational modifications will likely be missed. However, in the context of this study two PPIs were recovered which are dependent on a PTM. Further research is needed to see if the interacting fragments line up with those identified in the literature. Lastly, because of using short reads, the likely end point of the fragment is estimated based on the size-selected library length. However, this could be improved in the future by paired-end sequencing all fragments prior to the stitch PCR process to identify start and stop points for all fragments. It would be rare that two fragments would have exactly the same start point in a gene and so that would serve as an index to look-up its end point.

## Methods

### Reagents, strains, media, and plasmids

The human reference set was supplied in Gateway vectors^9^ (hsPRS-v2; hsRRS-v2). Each gene (Supplemental Table 1) was amplified individually using 1-5 ng DNA with primers sitting ~140 bp upstream and downstream of the gene. Minimal selection media, validation reporter cells and plasmid descriptions are as listed^10^. AVA-Seq plasmids are readily available from Addgene.org. All reagents are consistent with Andrews et al., 2019 unless stated otherwise.

### Library construction, screening and DNA sequencing

The method for library construction was similar to previously published^10^ with only slight modifications. Briefly, the 98 proteins used in this study were PCR amplified, quantified, and split into two pools of 20 nM each. PRS Batch 1 contained 39 proteins (22 PPIs) and Batch 2 contained 41 proteins (25 PPIs). Each Batch contained an additional 9 proteins that represented 6 RRS protein pairs (Supplemental Table 1, Supplemental Fig. 1). (Note: a few proteins are involved in both RRS and PRS interactions.) Each pool was sheared into ~500 bp fragments and processed as indicated in Andrews, et al. with the following changes^10^. Each sample included a positive (LGF2-Gal11p; 1:10^7^ dilution) and negative control (Gal11p-LGF2(fs); 1:10^7^ dilution) spiked in. Paired fragments in the pAVA vector were transformed into the reporter strain and 9 replicates were created. These 9 replicates were divided into 3 groups containing 3 replicates for 0, 2, and 5 mM 3-AT selection conditions and grown for 9 hours. DNA from the growth was extracted and libraries were generated using standard protocols. Samples were sequenced on an Illumina NovaSeq with paired 150bp reads according to the manufacturer’s recommended protocol.

### Primary Data Analysis

FASTQ files from the sequencers were analyzed as described previously^10^. Briefly, paired-sequence reads were translated in-frame with the appropriate fusion protein (lambda cI or RNAP) from the pAVA construct. Translated sequences were matched to a database of PRS proteins using the rapid protein aligner (DIAMOND)^20^ and the start point in the protein noted. Paired sequences which both matched in-frame and with a PRS protein were kept. In-frame fragment pairs were then collated and each time the exact pair with the same protein and start point was observed in separate read-pair, the count was incremented. Counts for each of the fragment pairs across all 9 replicates (3 × 0 mM, 3 × 2 mM, 3 × 5 mM 3-AT) were placed in a table for statistical analysis.

### Auto-activator removal

It was necessary to remove fragments that possibly interact with the system proteins RNAP (AD) and lambda cI (DBD). All in-frame fragments fused to RNAP were searched for interactions with more than 3 out-of-frame fragments fused to lambda cI. These are suspected to be due to the RNAP-fused fragments interacting directly with lambda cI rather than the fragment it is fused to. The same process was repeated for in-frame fragments fused to lambda cI that interacted with more than 3 RNAP-fused out-of-frame fragments. The value of 3 fragments or more was selected based on analysis of the average number of interactions a fragment had from the data. Only fragments that had more than 3 interactions were removed from the 2 mM 3-AT conditions analysis, the less stringent selective condition. Few fragments had more than 1 interaction in 5 mM 3-AT conditions that a trend could not be observed for excessive interactions with out-of-frame fragments.

### Scaling of data

Aside from scaling data based on varying read counts across the replicates, analysis of the raw data revealed multiple fragments pair that would rise to thresholds of interaction calling in the 5 mM but not in the less stringent 2 mM 3-AT conditions. Closer inspection of the data showed that standard RNA-seq algorithms called multiple negative interaction, that is, fragment pairs that decreased in proportion from 0 mM to 2 mM 3-AT. An analysis across the full data set revealed this as a trend, especially in 2 mM 3-AT where potentially thousands of fragment pairs might interact in the pool and, without sufficient sequencing depth, gives the impression that non-interacting pairs were decreasing in proportion. This was less of an issue in the more selective 5 mM 3-AT conditions where fewer fragment pairs interacted, causing sequencing reads to not be distributed across so many increasing pairs. Therefore, the read counts in 2 mM and 5 mM were scaled to have constant levels of non-interacting fragments across all replicates. To scale the data, fragments which had more than 10 counts per replicate were taken into consideration as having sufficient levels of sampling. Two separate distributions of fragments were observed when average values from 2 mM (or 5 mM) were compared to 0 mM 3-AT. Distributions centered below 1 (the value of counts is smaller in 2 mM or 5 mM than in 0 mM) were set as deriving from non-interacting fragments. For each library, the mean of the distribution of average values in 2 mM or 5mM with respect to 0 mM were taken as a reference levels and assigned values of 1. All counts values in 2 and 5 mM were scaled according to these factors.

### Statistical analysis

Growth in 2 mM or 5 mM 3-AT as detected by scaled read count values compared to 0 mM were considered as a signal of a potential PPI. Statistical significance of differential growth was evaluated from 3 replicates in each growth condition. Only those fragment pairs that had at least 10 CPM (counts per million) across all replicates were taken for differential growth analysis. The R package edgeR^21^ was utilized to identify fragment pairs that showed a statistical increase in selective conditions (2 mM or 5 mM 3-AT) over background (0 mM 3-AT). Internally, edgeR performs normalization of the counts values to adapt for varying sequencing depths as represented by differing library sizes. A negative-binomial model is fitted to determine differential growth using the Fisher’s exact test for significance testing, which computes *p* value and the adjusted *p* values (FDR) for each protein fragment pair. Upon further analysis, fragment pairs which had logFC > 1 and FDR < 0.1 in the presence of 3-AT when compared with 0 mM 3-AT were considered as possible interactions.

### Interaction Filtering

For the test of binary interactions, no filtering was applied beyond the test of statistical significance to better mimic a true binary test condition. For all-vs-all analysis, more stringent filters were applied for the removal of interactions with low support. The all-vs-all analysis required the following to report a protein-protein interaction: multiple fragments in either orientation, logFC > 1 and FDR < 0.1, or 1 fragment in either orientation with a logFC > 3 and an FDR < 0.01.

### Analysis of interaction space coverage

Fig. 2 denotes the percentage of the PRS protein-protein space covered by the screening method (RRS proteins were processed separately, Supplemental Fig. 1 and 2). Since AVA-Seq works with fragments the total space between two proteins as a matrix of dimensions *m × n* was considered, where *m* represents the length of the first protein in amino acids and *n* represents the length of the other protein in amino acids. Whenever fragments from a pair were tested, the part of the matrix corresponding to the amino acid area would be considered covered. In the case of complete protein-protein space coverage there would be enough fragments from both proteins to cover the space in the whole matrix. Otherwise, the corresponding percentage of the covered matrix would be considered as a percentage of the tested space between those two proteins. Data were then plotted according to the orientation of the fragments (AD or DBD associated).

## Acknowledgements

This research was supported by funding from Qatar Foundation to Weill Cornell Medicine in Qatar in the form of the BMRP2 grant. We thank the members of the WCM-Q Genomics Core for preparation of DNA libraries and data collection. We appreciate fruitful discussion and comments provided by Dr. Marc Vidal during the preparation of this manuscript and for providing access to the hsPRS-v2 library used in this study.

## Data availability

Sequences were deposited to the Sequence Read Archive of NCBI under the accession XXXXX.

## Author Contributions

S.S.-R. and J.A.M. conceived the idea and designed the study; S.S.-R. and N.M.A. performed experiments; S.S.-R., N.M.A. and Y.A.M. collected the data; S.S.-R., J.A., N.M.A., D.E.H. and J.A.M. analyzed the data; S.S.-R., D.E.H. and J.A.M. provided critical insight; S.S.-R., J.A., D.E.H. and J.A.M. wrote the manuscript.

## Supplementary Information

**Supplementary Table 1:** Breakdown of individual PPI taken from the hsPRS-v2 or hsRRS-v2. The protein interaction pair numbering is according to Choi, et al.^9^. For Batch 1 and Batch 2 the PRS proteins are listed first followed by the 6 RRS protein pairs directly below. Note there are several proteins which are in multiple protein interactions and a few proteins which are in both the positive interacting (PRS) and negative interaction (RRS) pairs.

| Batch 1 PRS |  |  | Batch 2 PRS |  |  |
| --- | --- | --- | --- | --- | --- |
| PAIR # | PROTEIN X | PROTEIN Y | PAIR # | PROTEIN X | PROTEIN Y |
| 2 | LMNA | LMNB1 | 10 | FANCA | FANCG |
| 3 | JUNB | BATF | 19 | FEN1 | PCNA |
| 5 | LCP2 | GRAP2 | 26 | CBLB | GRB2 |
| 6 | BAK1 | BCL2L1 | 27 | MCM2 | MCM3 |
| 7 | BAD | BCL2L1 | 31 | TP53 | UBE2I |
| 11 | PSMD4 | RAD23A | 32 | HIF1A | TP53 |
| 12 | LSM3 | LSM2 | 33 | GRB2 | VAV1 |
| 13 | MAD2L1 | MAD1L1 | 34 | SMAD1 | SMAD4 |
| 15 | MAFG | NFE2L1 | 37 | SMAD4 | DCP1A |
| 16 | PEX14 | PEX19 | 38 | CASP2 | CRADD |
| 17 | PEX19 | PEX16 | 39 | XIAP | CASP9 |
| 21 | GTF2F1 | GTF2F2 | 40 | NR3C1 | HSP90AA1 |
| 22 | IGF2 | IGFBP4 | 41 | ORC2L | ORC4L |
| 24 | PEX19 | PEX3 | 42 | RAC1 | ARFIP2 |
| 25 | LCP2 | NCK1 | 43 | ERBB3 | NRG1 |
| 28 | AKT1 | PDPK1 | 44 | CGA | CGB5 |
| 29 | RCC1 | RAN | 45 | ARF1 | ARFIP2 |
| 30 | NF2 | HGS | 47 | XIAP | CASP7 |
| 35 | CEBPG | FOS | 48 | IFIT1 | EIF3E |
| 46 | LMNA | RB1 | 49 | ORC2L | MCM10 |
| 50 | BDNF | NTF4 | 52 | XIAP | CASP3 |
| 51 | HDAC1 | ZBTB16 | 53 | GADD45A | PCNA |
|  |  |  | 55 | RIPK2 | NOD1 |
|  |  |  | 58 | PPP3CA | PPP3R1 |
|  |  |  | 59 | MCM2 | MCM5 |

| Batch 1 RRS |  |  | Batch 2 RRS |  |  |
| --- | --- | --- | --- | --- | --- |
| PAIR # | PROTEIN X | PROTEIN Y | PAIR # | PROTEIN X | PROTEIN Y |
| 89 | FAS | LSM3 | 85 | ERBB3 | C3orf38 |
| 129 | RCC1 | KLHL6 | 107 | MCM2 | PLXNA4 |
| 79 | COPB1 | HPCAL4 | 125 | PSMD5 | SLC22A15 |
| 80 | COPB1 | SLC39A14 | 126 | PSMD5 | SYCE1 |
| 92 | GALK1 | MCCC1 | 122 | PPP6C | ZNF350 |
| 93 | GCDH | ZCCHC9 | 123 | PROS1 | STK25 |

**Supplementary Table 2:**
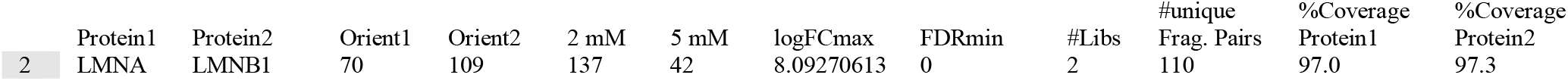
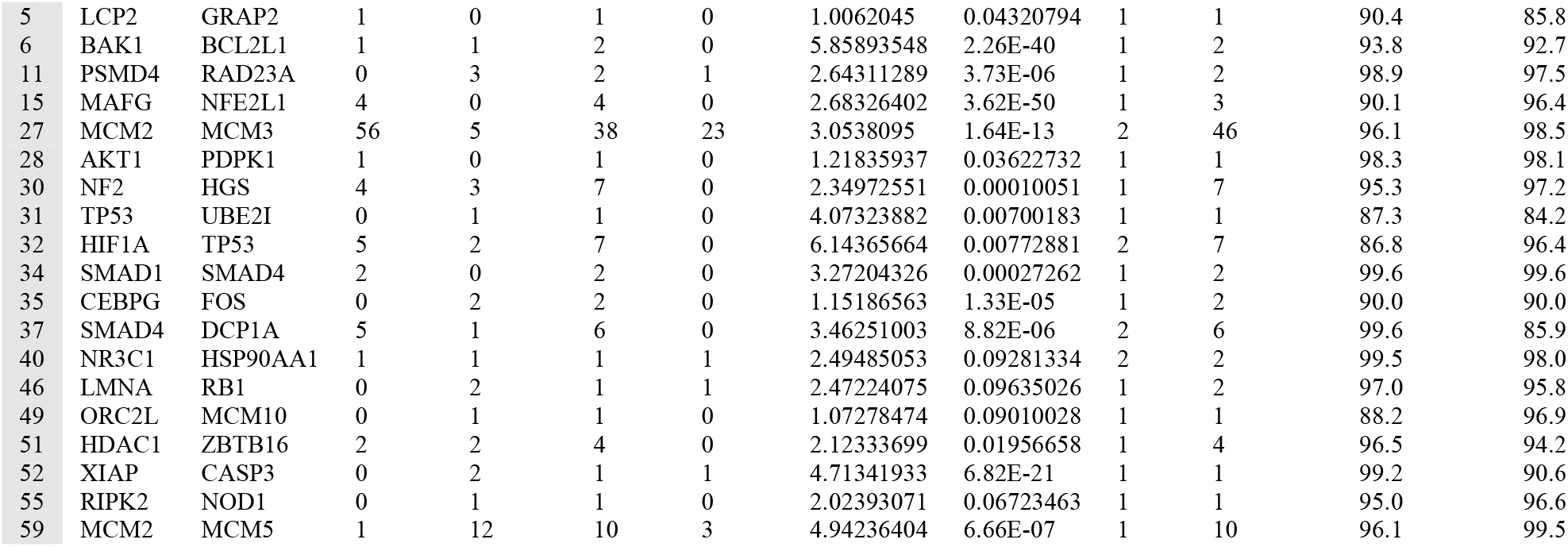
Observed binary interactions. A list of the 20 of 47 PPIs recovered from this study.

|  | Protein1 | Protein2 | Orient1 | Orient2 | 2 mM | 5 mM | logFCmax | FDRmin | #Libs | #unique<br>Frag. Pairs | %Coverage<br>Protein1 | %Coverage<br>Protein2 |
| --- | --- | --- | --- | --- | --- | --- | --- | --- | --- | --- | --- | --- |
| 2 | LMNA | LMNB1 | 70 | 109 | 137 | 42 | 8.09270613 | 0 | 2 | 110 | 97.0 | 97.3 |
| 5 | LCP2 | GRAP2 | 1 | 0 | 1 | 0 | 1.0062045 | 0.04320794 | 1 | 1 | 90.4 | 85.8 |
| 6 | BAK1 | BCL2L1 | 1 | 1 | 2 | 0 | 5.85893548 | 2.26E-40 | 1 | 2 | 93.8 | 92.7 |
| 11 | PSMD4 | RAD23A | 0 | 3 | 2 | 1 | 2.64311289 | 3.73E-06 | 1 | 2 | 98.9 | 97.5 |
| 15 | MAFG | NFE2L1 | 4 | 0 | 4 | 0 | 2.68326402 | 3.62E-50 | 1 | 3 | 90.1 | 96.4 |
| 27 | MCM2 | MCM3 | 56 | 5 | 38 | 23 | 3.0538095 | 1.64E-13 | 2 | 46 | 96.1 | 98.5 |
| 28 | AKT1 | PDPK1 | 1 | 0 | 1 | 0 | 1.21835937 | 0.03622732 | 1 | 1 | 98.3 | 98.1 |
| 30 | NF2 | HGS | 4 | 3 | 7 | 0 | 2.34972551 | 0.00010051 | 1 | 7 | 95.3 | 97.2 |
| 31 | TP53 | UBE2I | 0 | 1 | 1 | 0 | 4.07323882 | 0.00700183 | 1 | 1 | 87.3 | 84.2 |
| 32 | HIF1A | TP53 | 5 | 2 | 7 | 0 | 6.14365664 | 0.00772881 | 2 | 7 | 86.8 | 96.4 |
| 34 | SMAD1 | SMAD4 | 2 | 0 | 2 | 0 | 3.27204326 | 0.00027262 | 1 | 2 | 99.6 | 99.6 |
| 35 | CEBPG | FOS | 0 | 2 | 2 | 0 | 1.15186563 | 1.33E-05 | 1 | 2 | 90.0 | 90.0 |
| 37 | SMAD4 | DCP1A | 5 | 1 | 6 | 0 | 3.46251003 | 8.82E-06 | 2 | 6 | 99.6 | 85.9 |
| 40 | NR3C1 | HSP90AA1 | 1 | 1 | 1 | 1 | 2.49485053 | 0.09281334 | 2 | 2 | 99.5 | 98.0 |
| 46 | LMNA | RB1 | 0 | 2 | 1 | 1 | 2.47224075 | 0.09635026 | 1 | 2 | 97.0 | 95.8 |
| 49 | ORC2L | MCM10 | 0 | 1 | 1 | 0 | 1.07278474 | 0.09010028 | 1 | 1 | 88.2 | 96.9 |
| 51 | HDAC1 | ZBTB16 | 2 | 2 | 4 | 0 | 2.12333699 | 0.01956658 | 1 | 4 | 96.5 | 94.2 |
| 52 | XIAP | CASP3 | 0 | 2 | 1 | 1 | 4.71341933 | 6.82E-21 | 1 | 1 | 99.2 | 90.6 |
| 55 | RIPK2 | NOD1 | 0 | 1 | 1 | 0 | 2.02393071 | 0.06723463 | 1 | 1 | 95.0 | 96.6 |
| 59 | MCM2 | MCM5 | 1 | 12 | 10 | 3 | 4.94236404 | 6.66E-07 | 1 | 10 | 96.1 | 99.5 |

**Supplementary Table 3.** List of all significant interactions detected in this study. See attached.

**Supplementary Figure 1.**
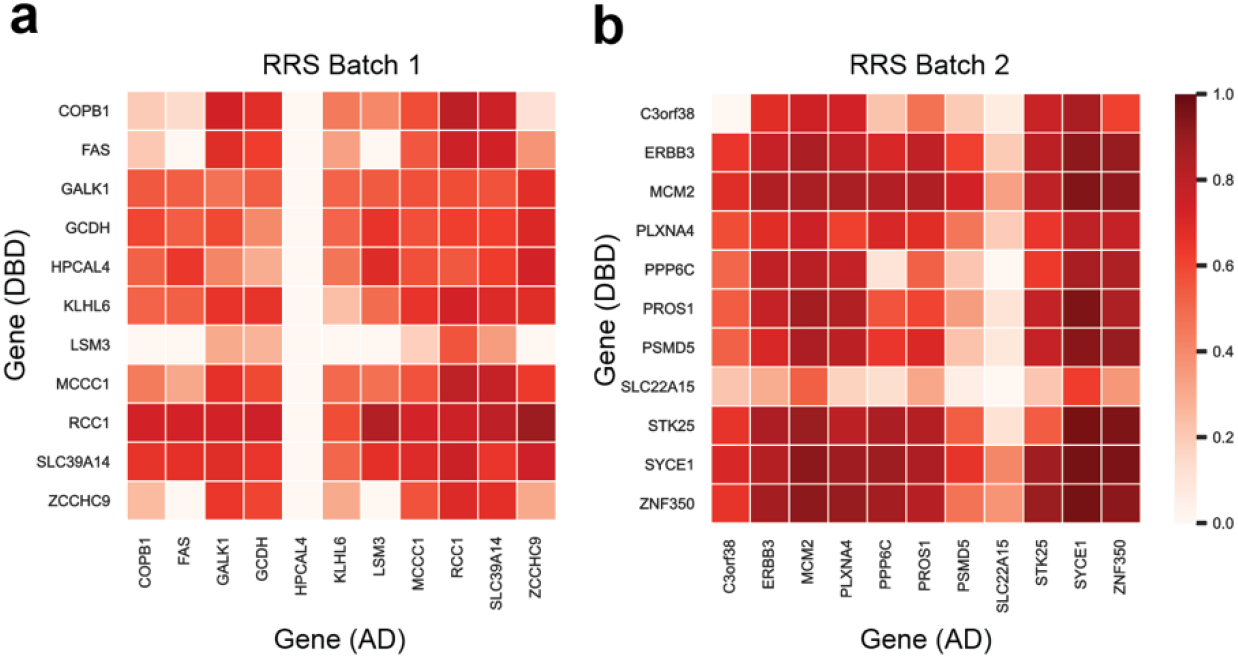
RRS coverage heatmap. Some of the proteins designated as an RRS protein pair are also in a PRS protein interaction.

**Supplementary Figure 2.**
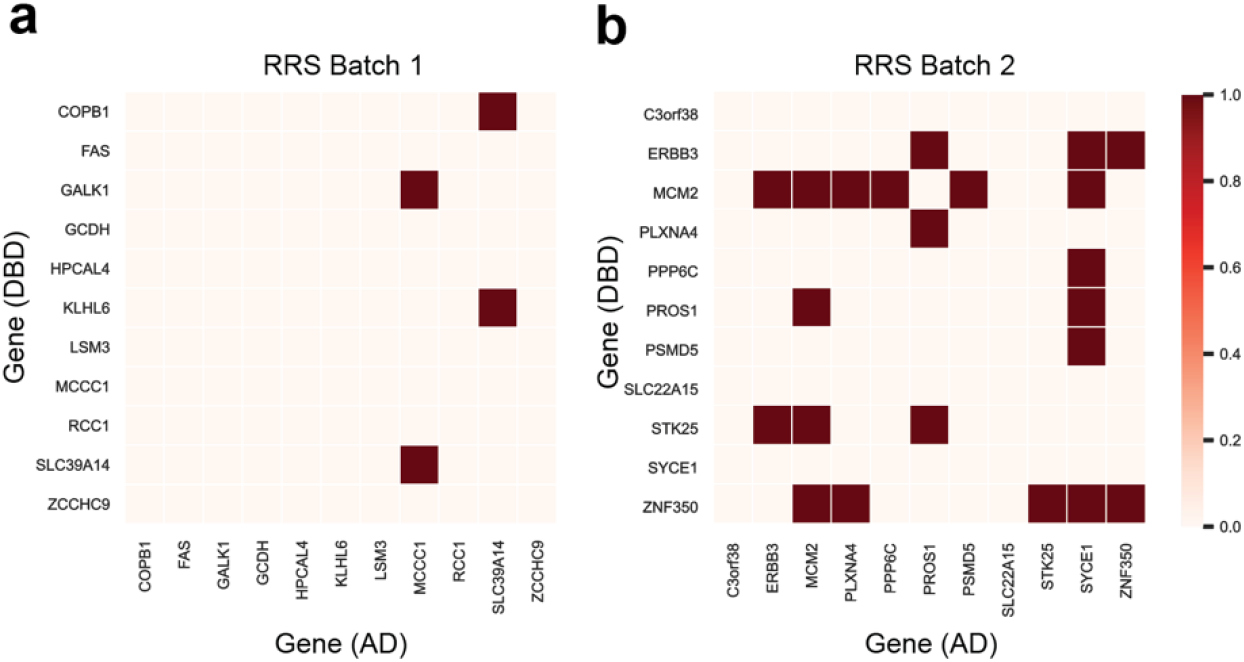
RRS interaction map. Summation of interactions detected in 2 mM and 5 mM 3-AT.

**Supplementary Figure 3.**
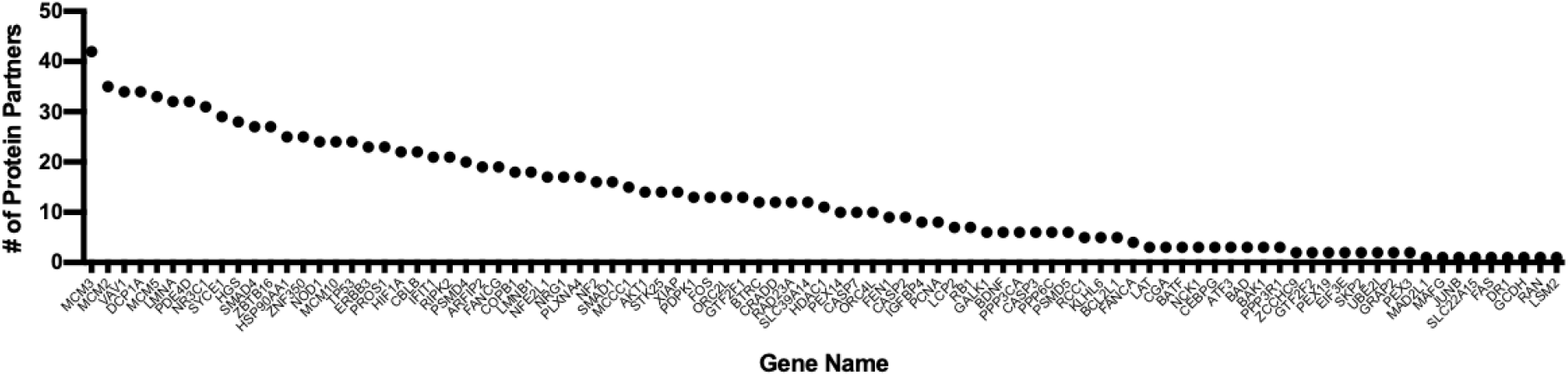
Number of interacting partners for the PRS proteins in the All-vs-All data.

**Supplementary Figure 4.**
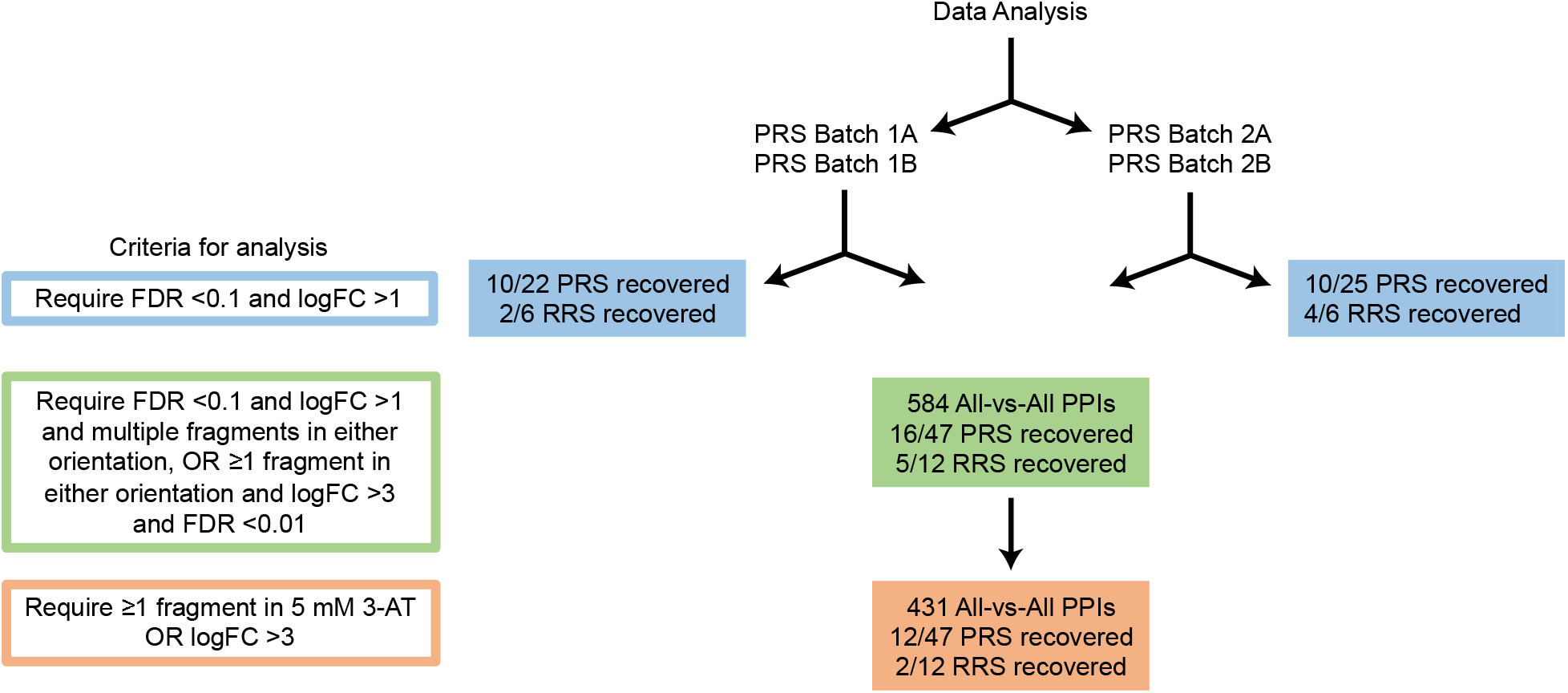
Graphical representation of criteria used for analysis along with number of recovered PPIs.

